# Zebrafish yolk syncytial nuclei migrate along a dynamic microtubule network

**DOI:** 10.1101/207795

**Authors:** Zhonghui Fei, Koeun Bae, Serge E. Parent, Katharine Goodwin, Guy Tanentzapf, Ashley E.E. Bruce

**Affiliations:** Department of Cell and Systems Biology, University of Toronto, Toronto ON M5S 3G5; Life Sciences Institute, Vancouver Campus, 2350 Health Sciences Mall, Vancouver, BC Canada V6T 1Z3

**Keywords:** zebrafish, epiboly, yolk syncytial layer, yolk syncytial nuclei, microtubule, LINC complex

## Abstract

In teleosts, the yolk syncytial layer is a multinucleate syncytium that functions as an extraembryonic signaling center to pattern the mesendoderm, coordinate morphogenesis and supply nutrients to the embryo. The zebrafish is an excellent system for studying this morphogenetically active tissue. The external yolk syncytial nuclei (e-YSN) undergo microtubule dependent epiboly movements that distribute the nuclei over the yolk. How e-YSN epiboly proceeds, and what role the yolk microtubule network plays is not understood but currently it is proposed that e-YSN are pulled vegetally as the microtubule network shortens from the vegetal pole. Data from our live imaging studies suggest that the yolk microtubule network is dismantled from the animal and vegetal regions and show that a region of stabilized microtubules forms before nuclear migration begins. e-YSN do not appear to be pulled vegetally but rather move along a dynamic microtubule network. We also show that overexpression of the KASH domain of Syne2a impairs e-YSN movement, implicating the LINC complex in e-YSN migration. This work provides new insights into the role of microtubules in morphogenesis of an extraembryonic tissue.

**Summary Statement:** Analysis of yolk syncytial nuclear migration during zebrafish epiboly reveals that nuclei migrate along and largely beneath a dynamically yolk microtubule network.

## Introduction

Embryonic development involves coordinated cell shape changes and movements to establish the adult body plan and developmental programs are inextricably linked to embryo architecture. In teleosts, the yolk syncytial layer (YSL) is a conserved and essential extraembryonic signaling center which contains transcriptionally active yolk syncytial nuclei (YSN). The YSL has numerous functions including induction and patterning of mesendoderm, coordination of epiboly movements and provision of nutrients to the embryo (Mizuno et al. 1999; Feldman et al. 1998; Ober & Schulte-Merker 1999; Rodaway et al. 1999; Gritsman et al. 2000; Koos & Ho 1998; Thomas 1968; Ho et al. 1999; Sirotkin et al. 2000; Fekany et al. 1999; Fekany-Lee et al. 2000; Chen & Kimelman 2000). In addition, the YSL undergoes surprisingly dynamic shape changes during development, making it an excellent system to gain new insights into morphogenesis of an extraembryonic tissue (Carvalho et al. 2009; D’Amico & Cooper 2001; Virta & Cooper 2011). YSL functions rely upon YSN transcription (Chen & Kimelman 2000; Xu et al. 2012) and a population of YSN undergo active epiboly movements which distributes the nuclei over the yolk surface (D’Amico & Cooper 2001; Carvalho et al. 2009). When the nuclei are not properly distributed, epiboly and patterning are adversely affected (Carvalho & Heisenberg 2010; Xu et al. 2012; Takesono et al. 2012).

The YSL forms as a result of meroblastic cleavages which generates the blastoderm on top of a large yolk cell (Carvalho & Heisenberg 2010; Kimmel & Law 1985; Trinkaus 1993). In zebrafish, incomplete early cleavages result in marginal blastomeres remaining open to the yolk cell and, around the time of the maternal-zygotic transition, marginal blastomeres release their cytoplasm and nuclei into the previously anuclear yolk cell to form the YSL. The YSL consists of the external-YSL (e-YSL) at the yolk-blastoderm exterior interface while the internal YSL (i-YSL) lies directly beneath the blastoderm (Kimmel & Law 1985). Yolk syncytial nuclei (YSN) are located in both regions and are referred to as e-YSN and i-YSN (Kimmel et al. 1995) (Fig. 1A). YSN undergo several mitotic divisions before they exit the cell cycle (Kane et al. 1992). YSN become enlarged and in some species, have been shown to become polyploid (Bachop and Schwartz, 1974).

**Figure 1.**
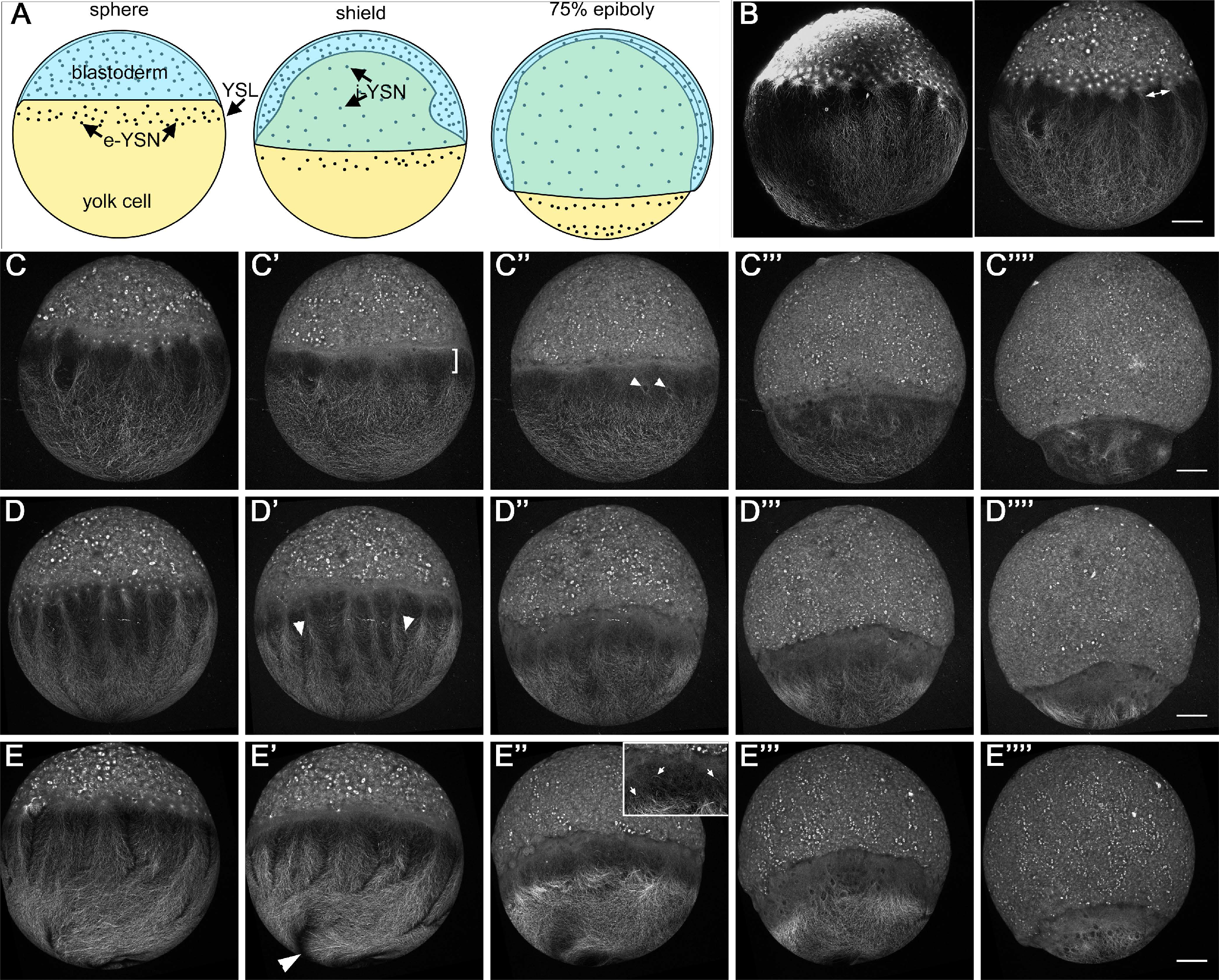
Changing yolk cell microtubule dynamics during epiboly. Panels are lateral views with the animal pole to the top. **(A)** Embryo schematics during epiboly, blastoderm, e-YSN, i-YSN, YSL and yolk cell indicated. **(B)** Sphere stage embryos. Left: alpha-tubulin antibody staining, right: Tg(XlEef1a1:dclk2DeltaK-GFP) embryo. Double headed arrow indicates e-YSN nucleating microtubule branches. **(C-E’’’)** Live confocal projections of 3 Tg(XlEef1a1:dclk2DeltaK-GFP) embryos from early to late epiboly (left to right). **(C’)** bracket marks the dim zone. **(C’’)** Arrowheads indicate e-YSN. **(D’’)** Arrowheads indicate gaps between microtubule branches. **(E’)** Arrowhead points to clearing region vegetally. **(E’’)** Inset shows magnified view of dim zone, arrowheads indicate microtubule fragments. Scale bar: 100 μm.

Epiboly is a major cell movement during teleost development. Epiboly involves the thinning and spreading of a multilayer of cells and the active motive force provided by the YSL is absolutely required for this process (Trinkaus 1963). During epiboly, the blastoderm and YSL spread down towards the vegetal pole to cover and enclose the yolk cell by 10 hpf (Fig. 1A). The blastoderm generates all embryonic tissues and it consists of an outer epithelial layer, the enveloping layer (EVL), which is tightly attached at its margin to the yolk cell and covers the underlying deep cells. In zebrafish, epiboly begins at 4.3 hours post-fertilization (hpf), following the cessation of YSN mitotic divisions and lineage specification of the EVL (Kimmel et al. 1995). The e-YSL has been shown to provide mechanical force necessary to drive epiboly via actomyosin motors (Behrndt et al. 2012; Cheng et al. 2004). There is also a distinct longitudinal microtubule array that is often disrupted in embryos with abnormal epiboly (reviewed in Lepage & Bruce 2010; Bruce 2016). The microtubule network is nucleated from the most vegetally positioned e-YSN in the YSL and oriented along the animal-vegetal axis with the microtubule plus ends extending into the yolk cytoplasmic layer (YCL), which is a thin layer of cytoplasm that surrounds the dense core of yolk granules (Strähle & Jesuthasan 1993; Solnica-Krezel & Driever 1994). During epiboly, the microtubule network shortens as the YCL is gradually replaced by the YSL (Solnica-Krezel & Driever 1994).

The dramatic morphogenetic changes in the YSL and its role in promoting epiboly have been recognized for some time, but much remains to be learned about the mechanisms that drive yolk morphogenesis. The yolk cell microtubules were first implicated in YSL epiboly in studies from Jesuthasan and Strahle (1993) and Solnica-Krezel and Driever (1994). Ultra-violet irradiation of cleavage stage embryos or treatment with the microtubule depolymerizing drug nocodazole resulted in delayed epiboly (Strähle & Jesuthasan 1993). Solnica-Krezel and Driever (1994) showed that treatment of late blastula stage embryos with nocodazole prevented the e-YSN from moving vegetally during epiboly. In contrast, vegetal movement of the blastoderm was not as strongly impaired, although epiboly movements were slowed. Taxol treatment, which stabilizes microtubules, slowed epiboly of the YSL and blastoderm to similar extents (Solnica-Krezel & Driever 1994).

Jesuthasan and Strahle (1993) proposed that the yolk microtubule network comprises part of the epiboly motor. They suggested that yolk microtubules act to expand the YSL via microtubule motors moving vegetally along the microtubules, and pulling the attached blastoderm along with it. In contrast, Solnica-Krezel and Driever (1994) found that epiboly of the e-YSN, but not of the blastoderm, was completely dependent upon the yolk microtubules. They postulated that the primary function of the network is to move the e-YSN vegetally during epiboly and that epiboly of the blastoderm is independent from epiboly of the e-YSN. They put forward several potential models in which different microtubule motors could provide pulling or pushing forces to move the e-YSN. One model was that, as the yolk microtubules shorten from the vegetally located plus ends, the e-YSN are pulled downwards (Solnica-Krezel & Driever 1994).

The positioning and movement of nuclei is important in a number of developmental contexts (Bone & Starr 2016). The linker of nucleoskeleton and cytoskeleton (LINC) complex has emerged as an important and conserved component of the nucleus that functions to connect structural elements in the nucleus to the cytoskeleton (Starr & Fridolfsson 2010). The complex consists of Sad1p/UNC-84 (SUN) and Klarsicht/ANC-1/Syne (KASH) proteins, located in the inner and outer nuclear membranes respectively. The LINC complex is capable of interacting with microtubules, centrosomes, F-actin, intermediate filaments and the microtubule motor proteins dynein and kinesin (Starr & Fridolfsson 2010; Chang et al. 2015). Well established examples of microtubule based nuclear movement include pronuclear fusion, muscle fiber development, and neuronal interkinetic nuclear migration (Bone & Starr 2016).

How e-YSN move towards the vegetal pole remains unclear. In addition, the dynamics of the yolk microtubule network have not been reported in detail during epiboly. Here we revisit these questions using live imaging and quantitative analyses. We show that the organization of the yolk cell microtubule network undergoes striking changes just prior to e-YSN movement, which have not previously been reported. We observed that e-YSN move vegetally through and largely beneath the microtubule network, in contrast to the current view. In addition, we show that the LINC complex appears to be involved in e-YSN epiboly. We present a new model for e-YSN movement that takes into account the observed changes in microtubule dynamics and proposes that microtubule motor proteins interact directly with e-YSN via the LINC complex to drive vegetal e-YSN movement.

## Results

### The yolk microtubule network undergoes dynamic changes during epiboly

The yolk microtubule network covers approximately 400 microns along the animal-vegetal (A-V) axis of the exposed multinucleate yolk cell at the start of epiboly (Kimmel et al. 1995). The yolk cell size is about ten times that of a typical cell, which may enable the formation of microtubule patterns that are not possible in smaller cells due to the differences in scale. To learn more about how e-YSN use microtubules for their movement, we examined the organization and dynamics of the network in live embryos. To accomplish this we used embryos from the previously characterized transgenic line Tg:(XlEef1a1:dclk2DeltaK-GFP) in which microtubules are indirectly labeled via binding of a microtubule associated protein fused to GFP (Sepich et al. 2011). For our analyses, we divided epiboly into early (dome to shield, 4.3-6 hpf), mid- (shield-75% epiboly, 6-8 hpf) and late (75% epiboly-bud, 8-10 hpf) stages (Fig. 1A).

At sphere stage, prior to epiboly initiation, the yolk cell microtubule network in both transgenic and alpha-tubulin stained embryos was nucleated from microtubule organizing centers associated with a subset of the most vegetally positioned e-YSN (Fig. 1B). Based on appearance, we refer to microtubules nucleated from an individual e-YSN as a ‘branch’ since they broadened as they extended vegetally (Fig. 1B). We also observed gaps between microtubule branches emanating from adjacent e-YSN (Fig. 1D’). Although the yolk microtubule network originated from the e-YSN, the widening of the branches vegetally suggested that vegetal microtubules were unlikely to be nucleated entirely from e-YSN associated microtubule organizing centers, given the size of the yolk cell and based on other analyses described below. This observation is consistent with work in *Xenopus* eggs showing that microtubules can be nucleated from existing microtubules and that interphase cytoplasm has the ability to support spontaneous microtubule growth (Ishihara et al. 2014).

To examine microtubule network dynamics, we generated low magnification confocal time-lapse movies of live transgenic embryos. In movies that captured the last YSN division during sphere stage, we observed that the division was accompanied by what resembled a wave of microtubule bundling followed by the re-establishment of the network from the e-YSN (Movie 1). This observation supports the idea that the e-YSN provide the polarity and the basic scaffold upon which the yolk microtubule network is built. We also note that this type of microtubule network is not observed in regularly sized cells, which would be encompassed within the width of a single microtubule branch.

The blastoderm began to spread vegetally shortly after the final YSN division. During early epiboly, a region of reduced fluorescence became increasingly apparent between the blastoderm margin and the vegetal microtubules in the YCL (Fig. 1C’, bracket). During mid-epiboly, this band of reduced fluorescence, which we refer to as the dim zone (DZ), moved vegetally ahead of the blastoderm. On close observation, it was apparent that microtubules were present in the DZ but they appeared to be more diffuse and microtubule fragments were rarely observed, suggesting a change in microtubule organization (Fig. 1E’’, inset). More intensely fluorescent, and potentially bundled, microtubules were apparent vegetal to the DZ. We also observed that in some dclk2DeltaK-GFP expressing embryos, microtubules cleared from the vegetal area of the yolk cell (Fig. 1E’, arrowhead). We postulated that this could result from depolymerization at the vegetal pole or from upward movement of the network towards the animal pole, or from a combination of the two. Around 60% epiboly, individual e-YSN began to move vegetally (Fig. 1C’’) which will be described below. As epiboly progressed the gaps between microtubule branches were less apparent (for example Fig. 1D’’’). During late epiboly, the DZ became less distinct as the network became disorganized (Fig. 1C’’’’, D’’’’, E’’’’). Our observations were consistent with reports that the microtubules shorten over the course of epiboly (Solnica-Krezel & Driever 1994) but the DZ in the upper region of the yolk suggested that depolymerization from the vegetal pole might not be the exclusive mechanism.

### The dim zone moves vegetally during epiboly

To confirm and quantify our observations, fluorescence intensity measurements and kymographs of dclk2DeltaK-GFP time-lapse movies were generated (Fig. 2A,B). We were able to define the DZ as a minimum between the blastoderm and the vegetal mass of microtubules in the yolk cell in A-V fluorescence profile plots taken from the center of the embryo (Fig. 2A). The fluorescence profiles revealed that the signal was high and relatively noisy in the blastoderm, as well as in the vegetal region of the yolk cell. By contrast, the DZ was characterized as a valley between the blastoderm and vegetal pole in which the fluorescence profile was smooth. These observations are consistent with the more diffuse organization of microtubules and an overall reduction in microtubules in the DZ. By tracking the DZ over time we observed that in all cases the DZ moved towards the vegetal pole (Fig. 2B). The mean speed of the DZ was constant at approximately 0.826 μm/min ± 0.146μm/min, until late epiboly when the DZ could no longer be reliably detected. Mean speeds for each embryo are given in Table 1. The speed of the DZ is roughly similar to the reported rate of 15% per hour for blastoderm epiboly from shield to bud stage (Kimmel et al. 1995), which is approximately equivalent to 0.75μm/min.

**Figure 2.**
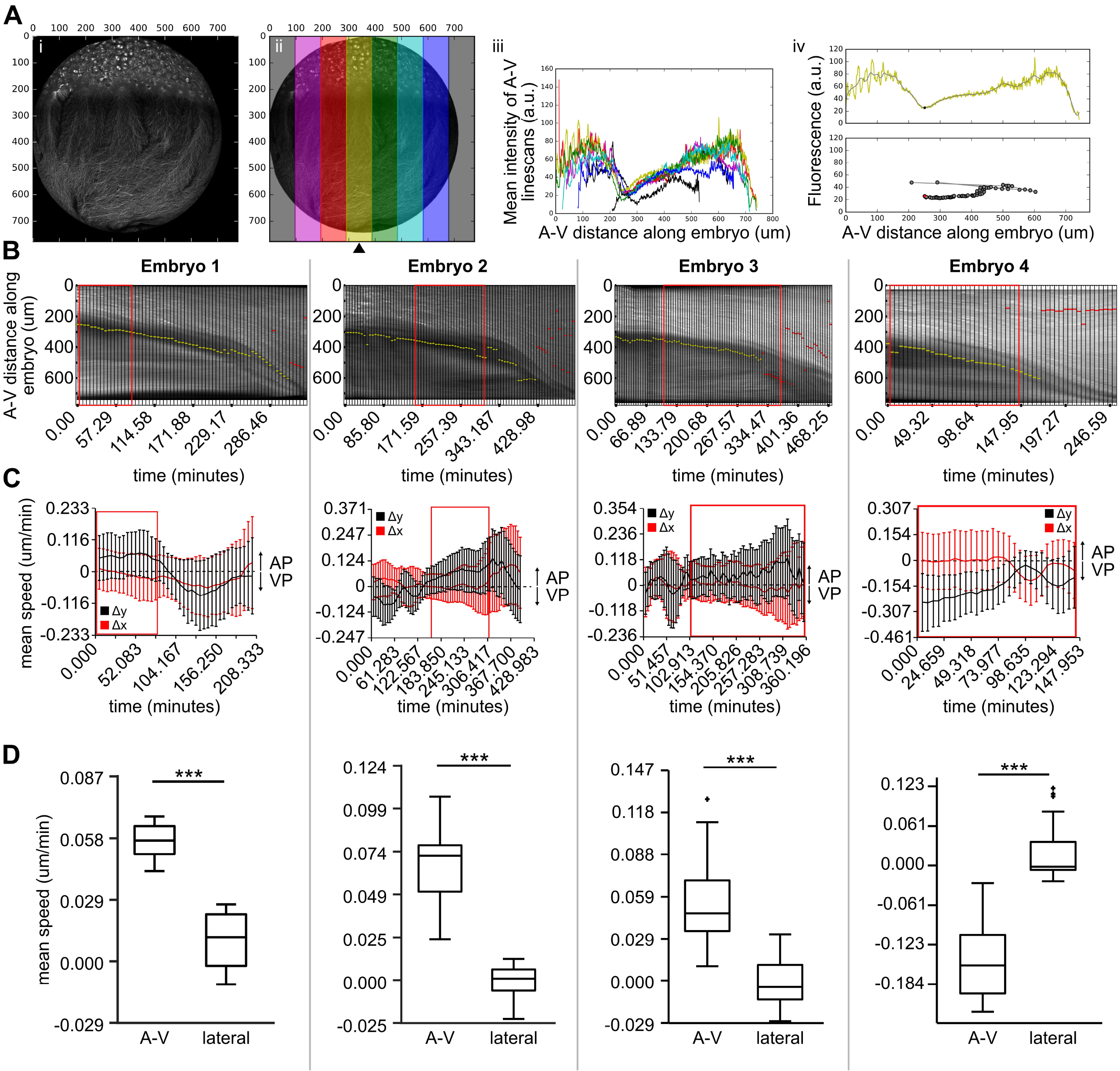
The dim zone moves vegetally and is preceded by animally moving microtubules from early to mid-epiboly. **(A)** Dim zone characterization. Masked time lapses showing only the embryo and no background (i) were separated in to 8 equally sized bins (ii). The mean across the lateral axis of each bin was taken and the resulting fluorescence profile was plotted (iii) and the centrally located yellow bin was used for subsequent analyses (arrowhead in “ii”). Minima present between the blastoderm and the vegetal microtubule network of the yolk were used to define the location of the dim zone (iv, top panel) and the position the minima was plotted over time (iv, bottom panel). **(B)** Kymographs of dim zone. Dim zone locations (i.e. minima) as determined by the mean yellow profile are denoted by horizontal yellow bars. Red bars denote dim zone locations that were false upon inspection and therefore removed from subsequent analyses. Red boxes mark early to mid-epiboly stages and correspond to the red boxes shown in (C). **(C)** PIV analyses of A-V directed and laterally directed flow of microtubules from early to mid-epiboly (red boxes). In the first 3 embryos, microtubule flow along the A-V axis (black lines) was directed animally, while lateral microtubule flow (red lines) was approximately zero. In the last embryo, microtubule flow was directed vegetally. Error bars show standard deviation. **(D)** Mean displacement of the microtubules per minute during early to mid-epiboly along A-V or lateral axes of the embryo. The data points represented in the plots are means for all vectors of a given time point. These sets of points for delta lateral and delta A-V were compared with 2-sided t-test using the matlab function ttest2, and *** indicates p<0.0001. Stages that red boxed regions correspond to: embryo 1: dome-60% epiboly; embryo 2: dome-75% epiboly; embryo 3: dome-60% epiboly; embryo 4: late sphere-65% epiboly.

**Table 1.**
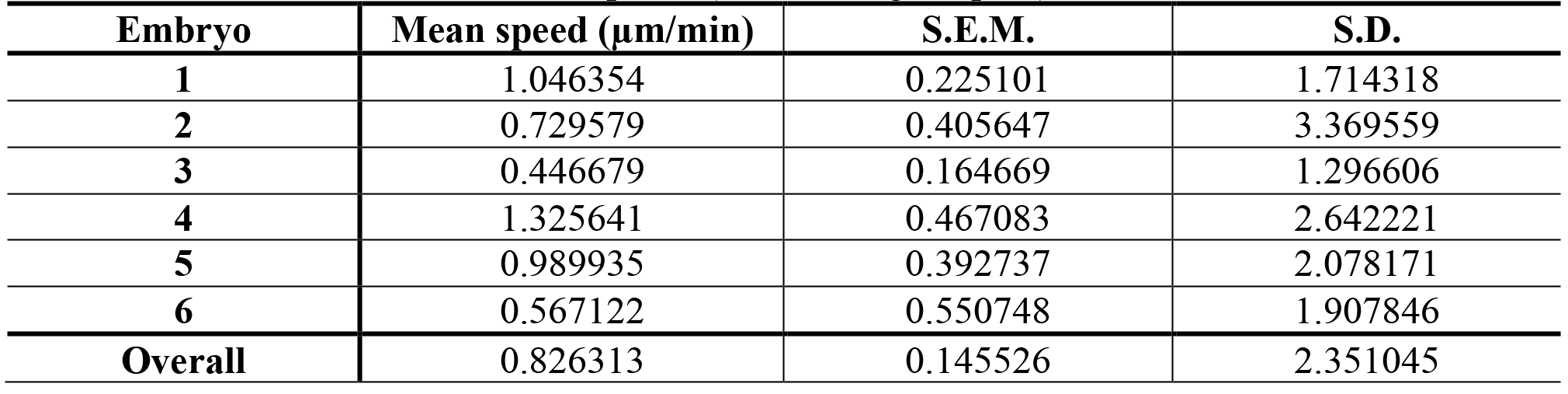
Mean dim zone movement speeds (towards vegetal pole).

Particle Image Velocimetry (PIV) analysis of four dclk2DeltaK-GFP time-lapse movies was done to further analyze microtubule movements (Fig. 2C,D). This analysis focused on the vegetal microtubules just below the DZ from early epiboly through the start of mid-epiboly. As expected, movement predominated along the A-V axis, rather than laterally. In embryos 1-3, there was greater mean displacement upwards towards the animal pole than downwards towards the vegetal pole (Fig. 2C red boxes, D). Embryo 4 differed in that there was greater mean displacement towards the vegetal pole than towards the animal pole throughout the time lapse (Fig. 2C,D). In all 4 embryos, the mean A-V speeds were an order of magnitude slower than the speed of the DZ. These findings suggest that the predominant event from early to mid-epiboly is the change in microtubule dynamics in the DZ and its vegetal progression. Microtubules within the DZ appeared to become more diffuse and the fluorescence was reduced. These observations suggest that the microtubule network is being dismantled, at least in part, from the top as the DZ moves vegetally. Below the DZ, fluorescence intensity was higher than within the DZ and microtubules were clearly visible. We hypothesized that this subset of microtubules might be more stable due to the accumulation of post-translational modifications.

### Detyrosinated tubulin is present in a subset of microtubules during mid-epiboly

To investigate the potential heterogeneity of the yolk cell network, we performed whole-mount antibody staining for detyrosinated tubulin, which is associated with longer-lived microtubules in vivo (Song & Brady 2014; Webster et al. 1987; Kreis 1987). Detyrosinated microtubules are typically present in cells, though often at very low levels, leading to the convention that detyrosinated microtubules are defined by detection over background using anti-detyrosinated tubulin antibodies (Bulinski & Gundersen 1991). Antibody staining in the yolk cell was technically challenging due to fixation, penetration and yolk trapping issues. Detyrosinated tubulin was detected by antibody staining in a subset of microtubules in the central region of the yolk of mid-epiboly stage embryos but was undetectable in sphere stage embryos (Fig. 3). Detyrosinated microtubules were located vegetally to the DZ and they did not extend to the vegetal pole. At 60% epiboly, migrating e-YSN could be seen about to enter this region (Fig. 3B, arrowhead). The antibody staining results were consistent with the idea that a subpopulation of stabilized detyrosinated microtubules is present during mid-epiboly outside the DZ.

**Figure 3.**
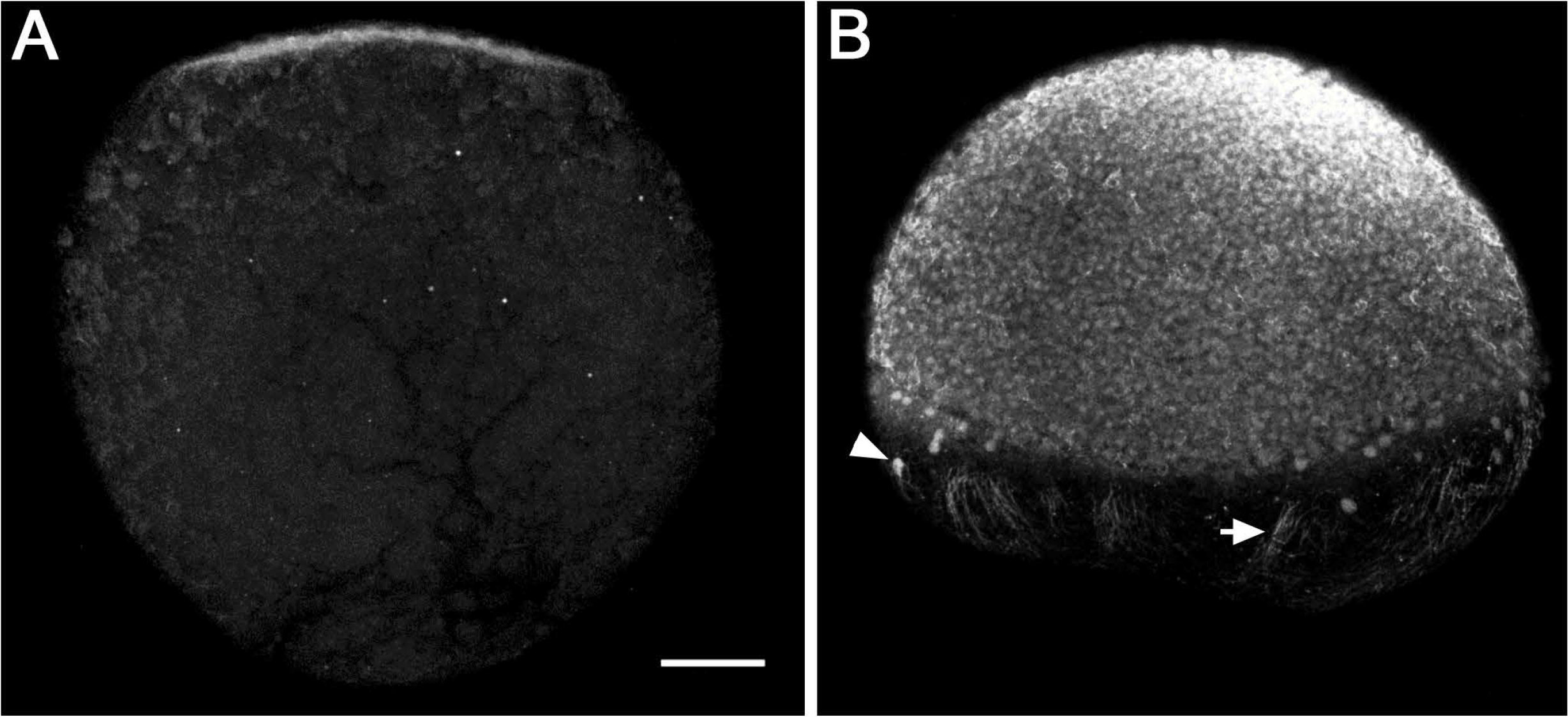
Detyrosinated microtubules are detected at mid-epiboly. Lateral views with the animal pole to the top of anti-detyrosinated tubulin stained embryos. **(A)** Sphere stage embryos showing absence of staining in the yolk cell, a small region of the blastoderm is visible at the top. **(B)** 60% epiboly stage embryo, detyrosinated tubulin is detected in the blastoderm and in the yolk cell (arrow). Arrowhead indicates e-YSN. Scale bar: 100 μm.

### EB3-GFP reveals widespread microtubule polymerization during early epiboly

To further characterize microtubule dynamics during epiboly, we investigated polymerization of the yolk cell microtubule network in live embryos by injecting *eb3-gfp* RNA into 1-cell stage embryos. EB3 is a microtubule plus end tracking protein that binds to actively growing microtubule plus ends and thus can provide information about the location and rate of microtubule growth as well as the polarity of the network (Stepanova et al. 2003). Embryos injected with *eb3-gfp* RNA were examined by confocal time-lapse microscopy from sphere to 85% epiboly. EB3-GFP fluorescent streaks (or comets) indicate active microtubule polymerization from the plus end.

Surprisingly, large numbers of EB3-GFP comets were visible throughout the yolk cell at sphere stage, indicating extensive microtubule growth (Fig. 4A, Movie 2). Some EB3-GFP comets clearly initiated at centrosomes associated with e-YSN while others could not be traced back to the e-YSN, providing support for the presence of non-centrosomal microtubules. EB3-GFP comets spread downwards towards the vegetal pole, consistent with the network having uniform polarity with microtubule plus ends extending vegetally (Solnica-Krezel & Driever 1994). Some comets curved laterally, consistent with morphology of the feather-like branches observed in dclk2DeltaK-GFP embryos. EB3-GFP comets were observed throughout early epiboly stages. Strikingly, during mid-epiboly the comets began to diminish and were largely undetectable in the YCL at late epiboly stages (Fig. 4A). Some EB3-GFP comets were still observed in the e-YSL, positioned close to the blastoderm, animal to the DZ. PIV analysis of a single plane time-lapse movie focused on the upper region of the yolk cell was consistent with our qualitative observations, showing that the predominant movement of EB3-GFP comets was along the A-V axis and directed towards the vegetal pole, with an average speed of 3.6 μm/min (Fig. 4B). The small amount of lateral movement might be explained by the feather-like shape of the microtubule branches.

**Figure 4.**
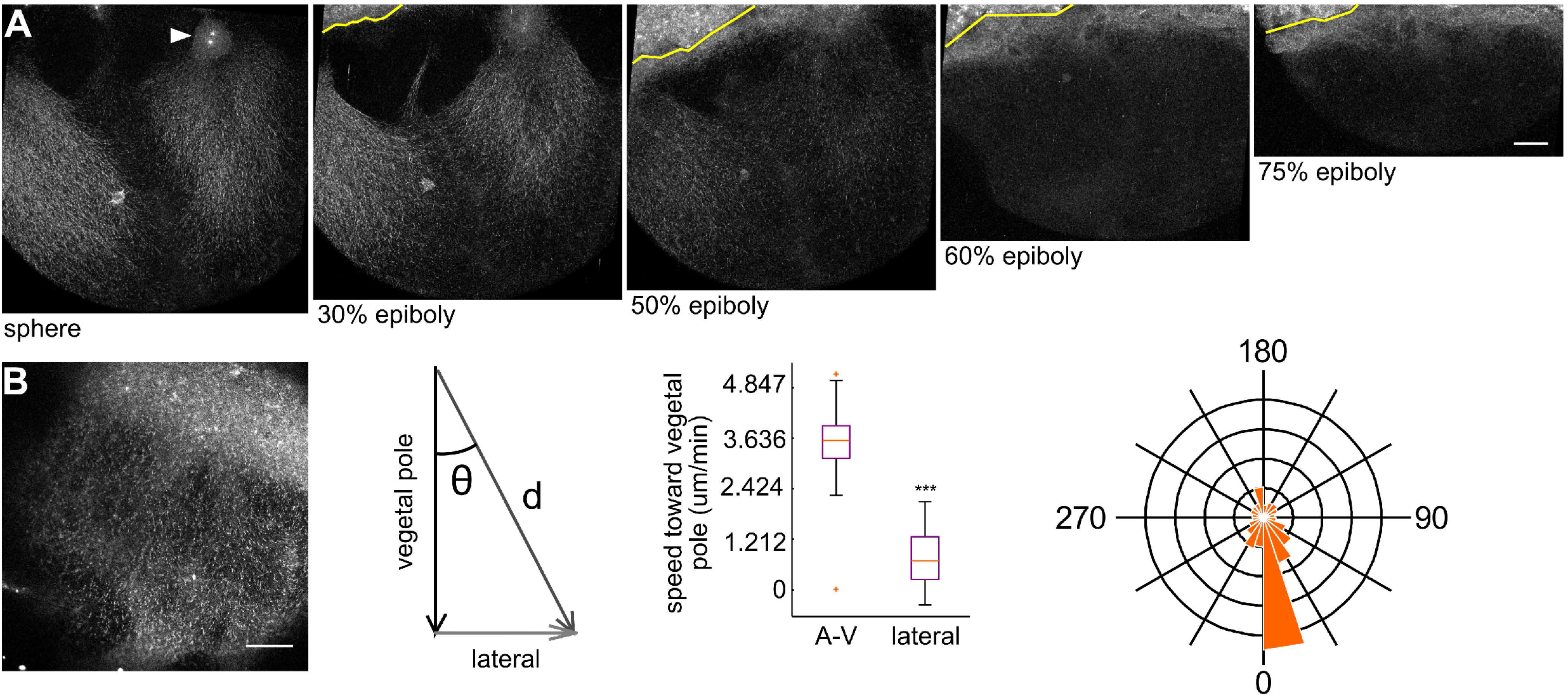
EB3-GFP reveals extensive microtubule polymerization during early epiboly. **(A)** Lateral confocal projections of wild-type embryos injected with *eb3-gfp* RNA, stages as indicated, animal pole up and slightly left. EB3-GFP fluorescent comets (visible as streaks) extend from the YSL into the YCL from sphere to 50% epiboly stage. By 60% epiboly comets were primarily confined to the e-YSL. Arrowhead indicates e-YSN centrosomes. Yellow lines mark boundary between the blastoderm and YSL. Scale bar: 50 μm. **(B)** PIV analysis of EB3-GFP fluorescent comet flow. Left panel shows still from single plane time-lapse movie analyzed. Blastoderm at the top right. Mean flow speed along A-V and lateral axes of the embryo measured as the mean of all vectors in a given time point. Positive flow along the A-V axis represents movement from the animal pole to the vegetal pole. These sets of points for lateral and A-V speed were compared with 2-sided t-test using the matlab function ttest2, and *** indicates p<0.0001. Right panel shows rose plot of PIV vector angles showing that vectors representing EB3-GFP movement are aligned with the A-V axis and directed vegetally.

To relate microtubule polymerization dynamics to nuclear movement, e-YSN in EB3-GFP movies were examined. Visible as non-fluorescent ovals, e-YSN could be seen to move out from regions where EB3-GFP puncta were being produced (Movie 3). As e-YSN moved vegetally EB3-GFP comets were visible behind them. Multiple e-YSN could be observed to move along microtubule branches being nucleated from stationary microtubule organizing centers. Currently it is not clear what causes the reduction in EB3-GFP comets and their confinement to the e-YSL. The reduction in puncta occurred during mid-epiboly, around the time that the DZ formed and detyrosinated microtubules were first detected. Intriguingly, these two events take place around that time that e-YSN begin to migrate. The temporal correlation between these events suggests that they might be linked and important for e-YSN movement.

### e-YSN move along and beneath the yolk microtubule network

The e-YSN start to move vegetally during mid-epiboly, after the formation of the shield (the zebrafish organizer) but what triggers the movement is not known (Solnica-Krezel & Driever 1994). We identified changes in the yolk microtubule network during mid-epiboly, before the e-YSN start to migrate. We postulated that initiation of e-YSN movement could be related to these changes. Thus, we sought to understand the relationship between the e-YSN and the yolk microtubules in more detail.

In low power time lapse movies of Tg:(dclk2DeltaK-GFP) and Tg:(XlEefla1:GFP-tuba8l) embryos, migrating e-YSN were visible as non-fluorescent ovals surrounded by fluorescent microtubules (Figs. 1C’’, 5A). Interestingly, as e-YSN began to migrate, they were often seen to move along the same trajectory. In a representative example, a single e-YSN moved vegetally and then shifted slightly medially, at which point a second e-YSN fell in line behind it and then a third e-YSN joined the line as if on a track (Fig. 5A). e-YSN often appeared to be linked, similar to previous reports of e-YSN chains connected by nuclear bridges (D’Amico & Cooper 2001). The strings of e-YSN were associated with microtubule branches extending from the YSL, confirming our observations that e-YSN move along EB3-GFP branches. Migrating e-YSN moved through the DZ, where the branches were less distinct, and then into the dense vegetal network of microtubules in the lower yolk.

**Figure 5.**
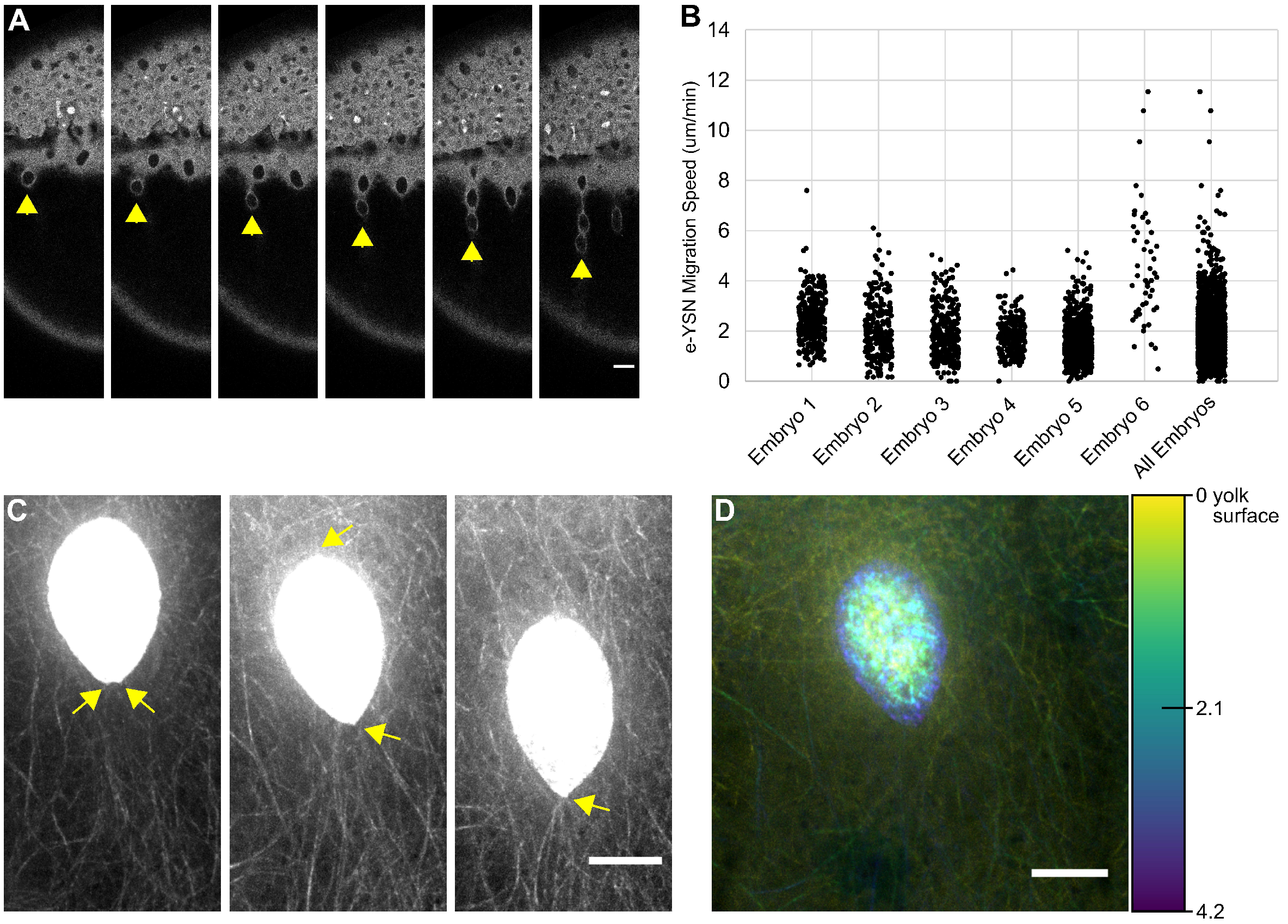
E-YSN move along and beneath the microtubule network. **(A-C)** Lateral views with the animal pole to the top. **(A)** Stills from a confocal time-lapse of a Tg:(XlEefla1:GFP-tuba81) embryo during mid-epiboly. Arrowheads indicate migrating e-YSN forming a chain. Scale bar: 25 μm. **(B)** e-YSN migration speeds from confocal-time lapse movies of 6 individual embryos and combined data. Embryos 1-5 were dclk2DeltaK-GFP transgenic embryos and embryo 6 was a GFP-tuba81 transgenic embryo. The time-lapse for embryo 6 was considerably shorter than for the other embryos which may explain the broader speed distribution. **(C)** Stills from spinning disk confocal time-lapse of Tg:(XlEefla1:GFP-tuba81) embryo injected with *h2a-gfp* RNA. The e-YSN becomes elongated and leading tip of e-YSN becomes pointed during migration (yellow arrows). Scale bar: 10 μm. **(D)** Depth coded projection from spinning disk confocal of Tg:(XlEefla1:GFP-tuba81) embryo injected with *h2a-gfp* RNA. The e-YSN is positioned largely beneath the microtubule network. Scale bar: 10 μm.

e-YSN speed was determined using 2D confocal projections of the time-lapse movies. e-YSN moved on average at approximately 1.936 ± 0.025 μm/min (see Table 2 for mean speeds per embryo; Table S1 mean speeds per nuclei). Nuclei did not appear to move at a uniform speed (Fig. 5B) but rather exhibited slow vegetal-ward movement punctuated by bursts of increased speed. These bursts of speed did not occur simultaneously, consistent with each e-YSN moving independently. Interestingly, the average speed was faster than blastoderm and DZ epiboly supporting the proposal that e-YSN epiboly is independent from blastoderm epiboly (Solnica-Krezel & Driever 1994).

**Table 2.**
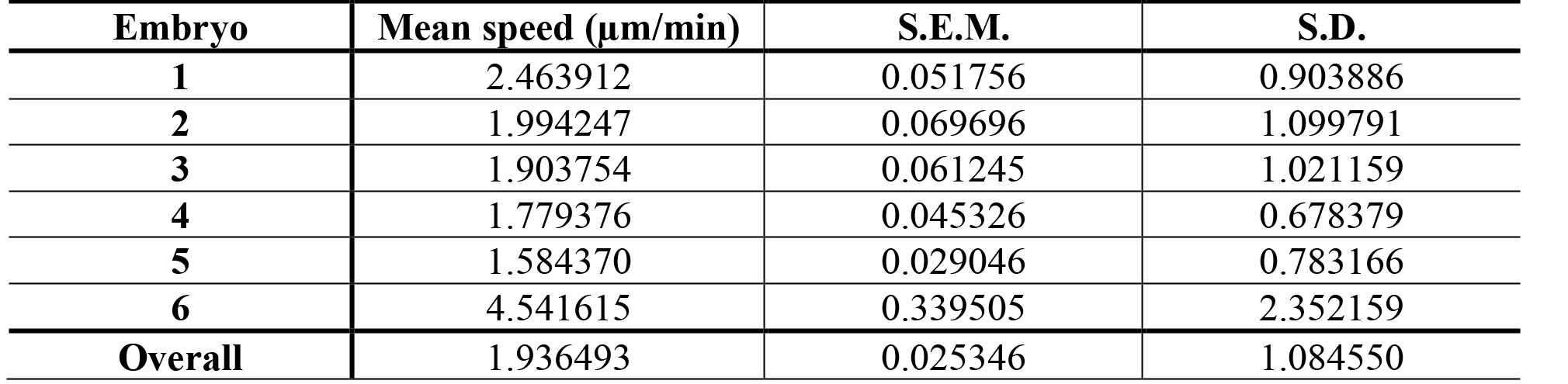
Mean e-YSN migration speeds per embryo (towards vegetal pole).

| **Embryo** | **Mean speed (µm/min)** | **S.E.M.** | **S.D.** |
| --- | --- | --- | --- |
| **1** | 2.463912 | 0.051756 | 0.903886 |
| **2** | 1.994247 | 0.069696 | 1.099791 |
| **3** | 1.903754 | 0.061245 | 1.021159 |
| **4** | 1.779376 | 0.045326 | 0.678379 |
| **5** | 1.584370 | 0.029046 | 0.783166 |
| **6** | 4.541615 | 0.339505 | 2.352159 |
| **Overall** | 1.936493 | 0.025346 | 1.084550 |

e-YSN movement was further examined using spinning disk confocal time-lapse microscopy of Tuba8l-GFP expressing embryos in which nuclei were fluorescently labeled with H2A-GFP. Consistent with Solnica-Krezel and Driever (1994) and our low power time-lapse movies, as the e-YSN began to move they typically underwent a shape change from round to elongate with the pointed end indicating the direction of movement (Fig. 5C). This shape change was most prominent around 60% epiboly, when the movement initiated. As the nuclei moved they exhibited small bulges and contractions on their surface (Movie 4). We confirmed that nuclear movements were continuous with short bursts of faster movement and migrating e-YSN were not observed to move backwards. Nuclei moved within individual microtubule branches and were not seen to cross the gap between branches. To understand the 3-dimensional relationship between the yolk microtubules and the e-YSN, we inspected Z-stacks from spinning disk confocal movies, and observed that the bulk of the microtubule network was more superficially located than the e-YSN (Fig. 5D). In the deepest e-YSN focal plane, microtubules were visible around the nuclei but were otherwise sparse compared to more superficial planes.

### Over-expression of a dominant-negative KASH construct disrupts e-YSN movement

There are several known mechanism whereby microtubules mediate nuclear migration (Gundersen & Worman 2013). In large cells, microtubules can exert pulling forces on centrosomes, which often involves cortically anchored dynein. Another method, common during developmental processes and exemplified by female pronuclear migration, involves nuclear envelope associated motor proteins ‘walking’ the nucleus down the microtubules (Gundersen & Worman 2013). Given our observation that the e-YSN move past and beneath the yolk microtubule network, we hypothesized that motor proteins move the e-YSN by directly associating with them. In addition, the formation of the DZ and the observation that e-YSN move through this region appears incompatible with vegetally anchored motor proteins pulling the e-YSN down the length of the yolk cell. We explored the possibility that the LINC complex, which is known to interact with microtubules and microtubule motor proteins (Starr & Fridolfsson 2010), was involved in e-YSN migration. Work in other systems, including the zebrafish retina, showed that overexpression of the KASH domain alone can impair nuclear movement by acting in a dominant-negative fashion to disrupt interactions between the LINC complex and cytoskeletal components or motor proteins (Tsujikawa et al. 2007; Grady et al. 2005).

To test the potential role of the LINC complex in e-YSN nuclear movement, we overexpressed the KASH domain of zebrafish Syne2a (C-syne2a) (Tsujikawa et al. 2007). Embryos were injected at the 1-cell stage with a mixture of *c-syne2a* and *h2a-gfp* RNA or with *h2a-gfp* RNA alone as a control. Confocal time-lapse microscopy was performed on injected embryos during mid-epiboly stages. In control embryos, e-YSN elongated in the direction of movement (Fig. 6 cell #1, Movie 5) as they moved towards the vegetal pole, as described above. In *c-syne2a* injected embryos, epiboly was overtly normal, however there were defects in the appearance and behavior of the e-YSN. The e-YSN did not become elongated to point in the direction of movement but were more globular in shape. Furthermore, instead of moving vegetally some e-YSN in *c-syne2a* injected embryos rotated sideways such that their movement was perpendicular to the A-V axis (Fig. 6 cells #2 and #3, Movie 6). Other e-YSN moved animally and some were overrun by the advancing blastoderm margin (Fig. 6 cell #1). These behaviors were not observed in control embryos. In *c-syne2a* injected embryos, most e-YSN were still carried vegetally, though in a less directed manner and we hypothesize that this movement is passive as a result of expansion of the YSL. These results suggest that SUN-KASH proteins are involved in directed e-YSN movement.

**Figure 6.**
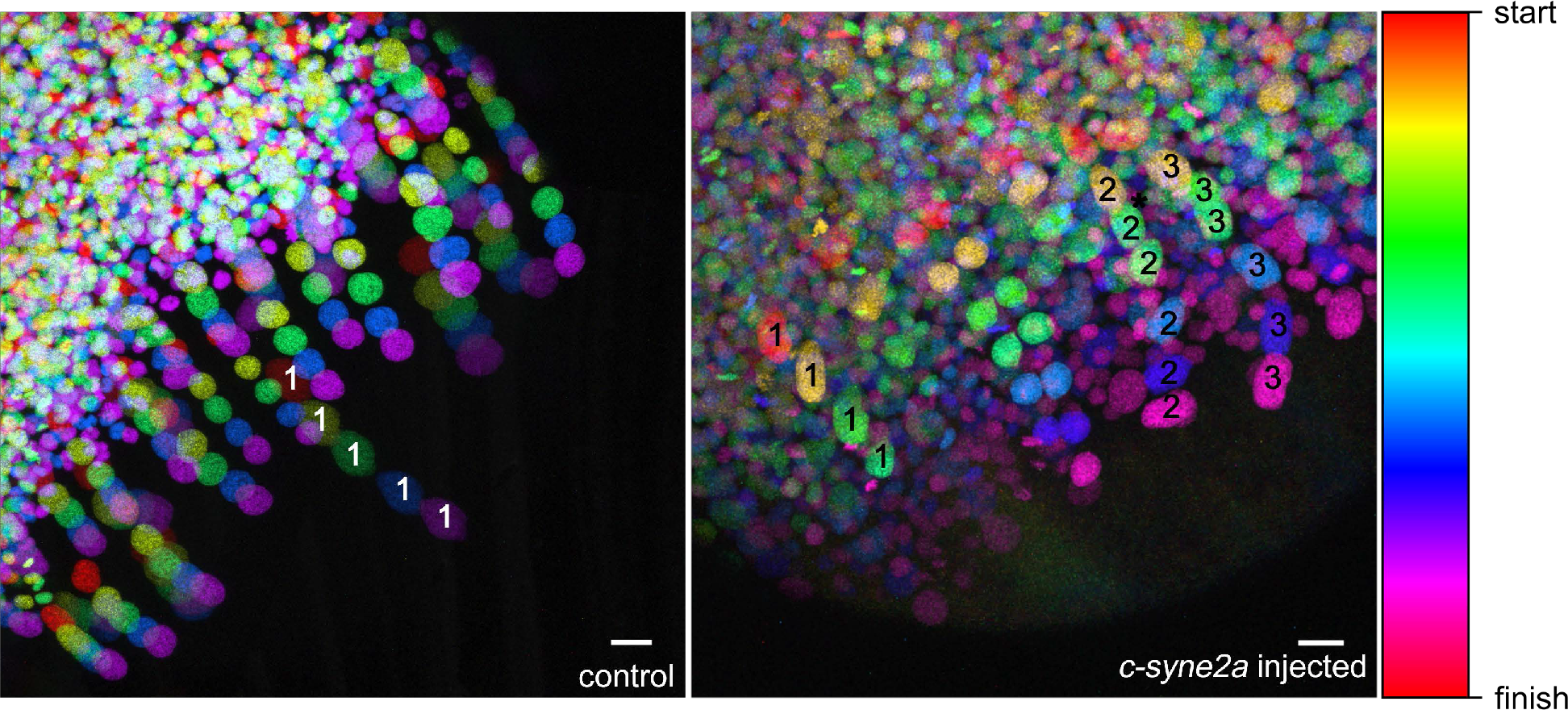
Overexpression of *c-syne2a* disrupts e-YSN migration. Temporal color coding of selected frames from confocal time lapse movies of *h2a-gfp* injected control and *h2a-gfp* plus *c-syne2a* injected embryos. Typical migrating e-YSN labeled with #1 in control embryo. In the *c-syne2a* injected embryo: cell #1 is overrun by the blastoderm, cell #2 turns perpendicular to the A-V axis and cell #3 starts to turn perpendicularly. Similar e-YSN behaviors were observed in *c-syne2a* injected embryos in 4 independent experiments.

We postulate that the LINC complex interacts directly with microtubule motors to move the e-YSN. Typically, centrosomes remain associated with nuclei as they migrate (Dupin & Etienne-Manneville 2011). Centrosomes can be located in front of the nucleus as it moves, with force transmitted to the nucleus via microtubules anchored at the centrosome (Solecki et al. 2004). In this scenario, dynein, a minus end directed motor, would be expected to drive nuclear movement in the yolk. However, the centrosome does not always lead the migration (Umeshima et al. 2007) and if this were the case, given the polarity of the yolk microtubule network, the plus end directed motor kinesin would be expected to be involved. To determine the location of the centrosome during e-YSN migration, we performed gamma-tubulin antibody staining. Gamma-tubulin was detected in association with e-YSN in the YSL at sphere and dome stages (not shown) but we were unable to detect gamma-tubulin in the yolk at later stages, due to background fluorescence. Our attempts to visualize the centrosome in live embryos during late epiboly by injecting RNA encoding Centrin-GFP or Gamma-tubulin-GFP were also unsuccessful. Thus, additional data are required to determine the position of the centrosome during nuclear migration as well as the motor protein involved.

## Discussion

The YSL is critically important for patterning and morphogenesis of the blastoderm. Previous work demonstrated that e-YSN undergo active microtubule dependent epiboly movements during gastrulation (Solnica-Krezel & Driever 1994; Carvalho et al. 2009; D’Amico & Cooper 2001). How epiboly of the e-YSN proceeds, and what role the yolk microtubule network performs during the process is not understood. Here we find that the e-YSN move along and largely beneath the cortical microtubule network. In addition, we identified changes in the structure and dynamics of the network that occur prior to e-YSN movement. Our results also implicate the LINC complex in e-YSN migration, confirming that microtubules are functionally involved.

### Yolk Microtubule Organization and Dynamics

The e-YSN nucleate initially non-overlapping microtubule branches which broaden and extend to the vegetal pole of the yolk cell. We propose that parts of the equatorial and vegetal network contain non-centrosomal microtubules. Non-centrosomal microtubules can be nucleated from golgi membranes, the nuclear envelop or from existing microtubules (Petry & Vale 2015; Lüders & Stearns 2007). We think it likely that vegetal microtubules are nucleated from existing microtubules, as reported for interphase *Xenopus* egg extracts (Ishihara et al. 2014). Although the zebrafish embryo is not as large as the *Xenopus* egg (750μm versus 1250μm), in both cases a yolk microtubule network forms over a distance that is much greater than a single cell. In addition, the smaller zebrafish embryo evolved from a large-egged ancestor similar to present day frogs (Cooper & Virta 2007), suggesting that mechanisms for generating the yolk microtubule network may be conserved. The *Xenopus* work also showed that microtubule based nucleation produces a network of parallel microtubules with uniform polarity (Petry et al. 2013), as is the case in the zebrafish yolk cell. Proof of this hypothesis will require the detection of gamma-tubulin at branched nucleation sites on yolk microtubules.

The yolk microtubule network has been assumed to be established prior to the start of epiboly and to be progressively shortened from the vegetal pole (Solnica-Krezel & Driever 1994). We were therefore surprised to see extensive EB3-GFP puncta throughout the yolk cell, which could reflect the non-centrosomal origin of some microtubules. During early epiboly, we also observed that the continuity between the upper and lower microtubule network began to diminish, as a region of reduced microtubule density appeared that we call the dim zone (DZ). Due to technical issues, we have so far been unable to simultaneously examine labeled EB3 and microtubules in live embryos which would allow us to understand the timing of these events in greater detail. The DZ was very obvious in dclk2DeltaK-GFP embryos and it was detected in GFP-tuba8l embryos. However, GFP-tuba8l embryos exhibit much lower levels of fluorescence than dclk2DeltaK-GFP embryos, which made the DZ more difficult to characterize.

After DZ formation, the structure of the microtubules in the DZ changed, although the details and molecular bases for these changes remains to be determined. Microtubules in the DZ appeared more diffuse and the overall reduction in fluorescence suggests that some were degraded, which was supported by time-lapse movies in which microtubule fragments entered the DZ and then lost their fluorescence. The DZ became more prominent during mid-epiboly stages as it moved as a wave front towards the vegetal pole as the microtubules vegetal to it shortened. One possibility is that the DZ represents the leading edge of YSL, which replaces the yolk cytoplasmic layer during epiboly. Towards the end of epiboly, the network became disorganized and the DZ was no longer apparent.

Microtubule dynamics are controlled by a number of factors including the tubulin isoforms being expressed (Panda et al. 1994), tubulin post-translational modifications (Janke & Chloë Bulinski 2011), and the activity of different microtubule associated proteins (MAPs) and motors (Heald & Nogales 2002; van der Vaart et al. 2009). A possible cause for the appearance of the DZ could be due to cleavage of microtubules by the microtubule severing protein Katanin, which has previously been reported to play a role in YSL epiboly (Bruce & Sampath 2008). Katanin severing in the upper region of the yolk cell would generate microtubules with free minus ends which would be expected either to be depolymerized or stabilized (Akhmanova & Hoogenraad 2015). Microtubules could be stabilized by the minus end binding protein Camsap2a. Camsaps are a recently identified family of microtubule minus end binding proteins and in other systems, Camsap2 plays a critical role in stabilizing non-centrosomal microtubules (Akhmanova & Hoogenraad 2015). Expression of zebrafish *camsap2a* is first detected in the e-YSL at sphere stage (Xu et al. 2012; Hong et al. 2010) where it may have a similar role. Katanin activity is regulated in a variety of ways and Katanin has been shown to regulate Camsap activity (Akhmanova & Hoogenraad 2015; Bailey et al. 2015). We postulate that these regulatory mechanisms control the extent of the DZ. During mid-epiboly, we detected a population of detyrosinated microtubules just vegetal to the DZ. Detyrosinated microtubules are relatively long-lived and interestingly, it has been shown that Camsap2 preferentially interacts with detyrosinated microtubules in cultured U2OS cells (Jiang et al. 2014). Detyrosinated microtubules are known to pause their growth due to capping (Infante et al. 2000), which could also contribute to the reduction in EB3-GFP puncta we observed during mid-epiboly. The stabilization of a subset of microtubules, by Camsap or other MAPs, would then enable further stabilization via the accumulation of post-translational modifications (Janke & Chloë Bulinski 2011).

We propose that timed expression of MAPs could be involved in the observed changes in microtubule dynamics. At present, the only characterized MAP in the zebrafish yolk cell is Clip1a, a zebrafish CLIP-170 homolog (Weng et al. 2013). Work from these authors showed that the steroid pregnenolone is required for normal epiboly and is involved in yolk cell microtubule stabilization (Hsu et al. 2006). They subsequently showed that pregnenolone functions by binding to Clip1a which then stimulates microtubule polymerization (Weng et al. 2013). As Clip1a is maternally expressed and CLIP-170 has been shown to have a preference for tyrosinated tubulin, it might be involved in the establishment of the network, but this remains to be tested (Ikegami & Setou 2010; Hsu et al. 2006).

Another open question is why the microtubule network is dismantled from the top and bottom. One possibility we are currently pursuing is a potential connection between the DZ and the actin cytoskeleton. The DZ is near the actomyosin cable that constricts to close the blastopore during epiboly (Cheng et al. 2004; Behrndt et al. 2012). In some migratory cells, depolymerizing microtubules can stimulate actin contractility via RhoA, and RhoA can in turn stabilize microtubules (Takesono et al. 2010; Chang et al. 2008; Wojnacki et al. 2014; Palazzo et al. 2001). This raises the interesting possibility of cross regulatory interactions between the microtubule and microfilament networks in the yolk cell that might be important for epiboly. We found that microtubule dynamics change around the same time that the actomyosin band becomes active (Behrndt et al. 2012). In addition, we observed that the DZ moves at a similar rate as the blastoderm, and blastoderm movement is driven by the yolk actomyosin motors. An interaction between these two cytoskeletal networks could help explain why in many examples of defective epiboly, both actin and microtubules are disrupted (Lachnit et al. 2008; Lee 2014).

### e-YSN Migration

Before e-YSN begin migrating, the DZ forms, a subset of microtubules become detyrosinated and EB3-GFP puncta diminish. Initially, e-YSN move along microtubule branches emanating from microtubule organizing centers producing EB3-GFP puncta and not from regions in between. This suggests that e-YSN nucleate microtubule tracks for other e-YSN to migrate along. Our model is that motor proteins, recruited to the e-YSN by the LINC complex, transport the nuclei through and past the DZ towards the vegetal pole. Thus, a given e-YSN is not linked to one set of microtubules throughout its movement and e-YSN appear to move through and largely beneath the bulk of the microtubules. We also saw that in some case microtubules moved slowly animally, in the opposite direction from migrating nuclei. Taken all together, these data do not support a model in which e-YSN are pulled by microtubule shortening from the vegetal pole.

The LINC complex is implicated in nuclear movement in many systems, and in keeping with this, we find that expression of a dominant-negative KASH domain construct impaired directional movement of e-YSN. Although our data do not allow us to define the motor protein involved, we suggest kinesin-1 as a likely candidate. In other systems the plus end directed microtubule motor kinesin-1 has high affinity for detyrosinated microtubules (Ikegami & Setou 2010). Interestingly, recent work using cultured fibroblasts and neurons demonstrated a novel function for kinesin-1 in promoting the formation of detyrosinated microtubules (Yasuda et al. 2017). Zebrafish kinesin-1, Kif5Ba, is maternally deposited and expressed throughout development (Campbell & Marlow 2013). We propose that detyrosination of the network facilitates kinesin-1 interaction with microtubules and that kinesin-1 is recruited to e-YSN via the LINC complex. Additional functional studies, as well as determining the location of the centrosome during e-YSN migration, will help clarify the mechanism of nuclear movement. To date the only KASH domain protein characterized during early zebrafish development is Lymphoid restricted membrane protein, which is involved in centrosome-nucleus attachment in the zygote (Lindeman & Pelegri 2012). The LINC complex has been implicated in nuclear migration in the zebrafish retina (Tsujikawa et al. 2007) and several uncharacterized LINC complex genes are present in the genome.

Our work supports previous reports on nuclear migration in the *C. elegans* epidermis, the hypodermis. The hypodermis undergoes epiboly during ventral enclosure and is essential for axis elongation (Williams-Masson et al. 1998; Williams-Masson et al. 1997). Hypodermis morphogenesis involves cell intercalation and microtubule dependent nuclear migration and the hyp7 hypodermal precursor cells are used as a model for nuclear migration (Williams-Masson et al. 1998). The KASH domain protein UNC-83 recruits kinesin-1 to the nuclear envelop to drive nuclear migration along microtubules (Meyerzon et al. 2009). Microtubules are nucleated from nuclei that are initially positioned at the immobile side of the cell such that a parallel array of microtubules forms with the plus ends oriented toward the intercalating side of the cell (Meyerzon et al. 2009). The network is thought to become non-centrosomal since nuclei migrate in association with their centrosomes. However, the centrosome does not lead the migration, consistent with the role of the plus end directed motor kinesin-1 (Meyerzon et al. 2009). The similarities in this system to the zebrafish yolk cell suggest that this mode of nuclear migration may be evolutionarily conserved. Important differences include the size difference between *C. elegans* cells and the zebrafish yolk cell and the presence of microtubule branches in zebrafish.

Our findings are summarized and assembled into a time line in Fig. 7. As epiboly starts, the DZ becomes apparent, it then moves towards the vegetal pole during epiboly until the late epiboly when the entire microtubule network becomes disorganized. A subset of detyrosinated microtubules becomes detectable at mid-epiboly as the e-YSN begin to migrate towards the vegetal pole. E-YSN migrate more rapidly than the blastoderm and DZ, and their migration requires the LINC complex.

**Figure 7.**
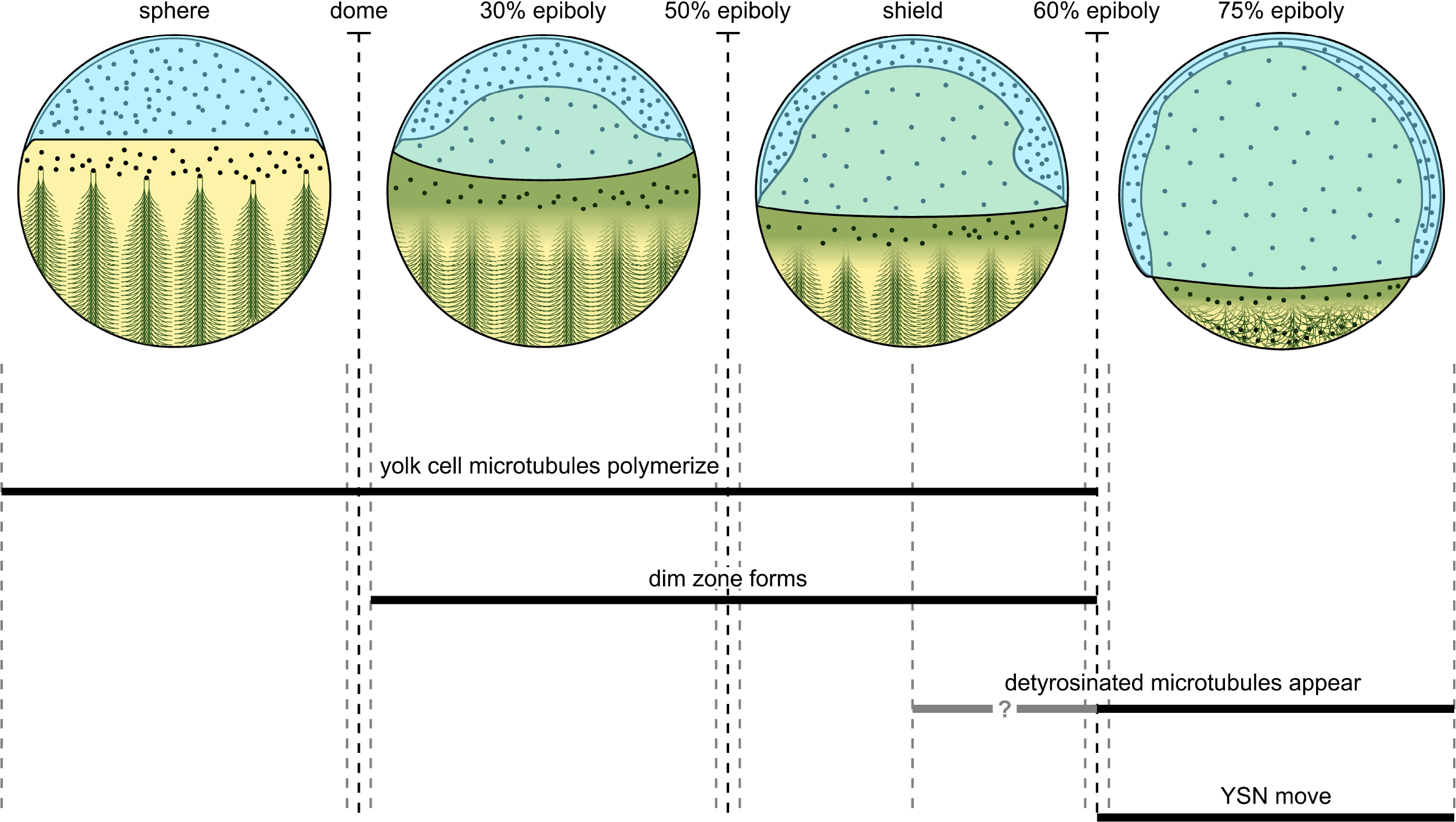
Model and Time-Line. Lateral views of schematic embryo with YSN and microtubules depicted, animal pole to the top and event timing below. Dashed black lines indicate stages and dashed grey lines indicate beginning and end of a given stage. bThe dim zone is represented by the green shaded area. Exactly when detyrosinated microtubules appear is unclear, as indicated by gray bar and question mark.

## Conclusions

The YSL is conserved in teleosts and is present in other organisms with meroblastic cleavage including the longnose gar, squid and chicken (Long & Ballard 2001; Wadeson & Crawford 2003; Arendt & Nübler-Jung 1999; Nagai et al. 2015). The critical signaling function of the YSL might explain why it is necessary for YSN to be distributed over the yolk surface during epiboly. During gastrulation different signals are sent to the dorsal and ventral sides of the blastoderm and gene expression in the YSL is temporally dynamic (Carvalho & Heisenberg 2010; Sun et al. 2014; Thisse & Thisse 2004). One proposal is that the YSL provides stabilizing signals that enhance the robustness of gastrulation and help maintain regional expression domains in the blastoderm as widespread cell rearrangements occur (Sun et al. 2014). Later developmental events also depend upon YSL signaling, such as heart morphogenesis (Trinh & Stainier 2004). The distribution of YSN is also likely to be important for the nutritive function of the yolk cell since, as lecithotrophs, zebrafish rely exclusively on the yolk for the first 5 days of development.

## Materials and methods

### Zebrafish strains

Zebrafish (*Danio rerio*) were maintained under standard conditions. AB, Tg(XlEef1a1:dclk2DeltaK-GFP) (gift from Marina Mione) (Sepich et al. 2011), and Tg (XlEef1a:eGFP-tuba8l) strains were used. Embryos were acquired from natural spawnings, maintained at 25-30°C and staged as described (Kimmel et al. 1995). Animals were treated in accordance with the policies of the University of Toronto Animal Care Committee.

### C-Syne2a construct

CDNA from 1 day post-fertilization embryos was synthesized using the Protoscript II 1^st^ Strand cDNA Synthesis kit (NEB) following the manufacturer’s instructions. The region of zebrafish *syn2a* encoding the KASH domain was PCR amplified from cDNA using Q5 high fidelity Taq polymerase (NEB) using the forward primer: 5’-CCACCATGCGCTCGTTCTTCTACCGTGT-3’ and reverse primer: 5’-TCATGTTGGAGGAGGGCCGT-3’. The PCR fragment was digested with EcoRI and ligated into EcoRI digested pCS2+ (Rupp et al. 1994). Orientation was confirmed by sequencing (TCAG DNA Sequencing Facility, Hospital for Sick Children).

### RNA Synthesis and Microinjection

To generate *h2a-gfp*, *eb3-gfp*, and *c-syne2a* sense RNA, NotI digested plasmids were in vitro transcribed using the SP6 mMESSAGE mMACHINE kit (Ambion). RNAs were purified with the MEGAclear kit (Ambion). Embryos were injected into the yolk of 1-cell stage embryos as described (Bruce et al. 2003). Doses of injected RNA were: *eb3-gfp* (110 pg), *h2a-gfp* (25 pg), and *c-syne2a* (50 pg).

### Generation of Tg:(XlEefla1:GFP-tuba8l) Transgenic Zebrafish

To generate an GFP-tubulin fusion construct, primers were designed to amplify the coding region of zebrafish *tubulin*, *alpha 2* (*tuba2*). The forward primer was 5’- ATGCGTGAGTGTATCTCCAT-3’ and the reverse primer was 5’-CTAATACTCCTCACCTTCCT-3’. RT-PCR was performed on shield stage cDNA and the PCR product was cloned into pGEM-T Easy (Promega). Sequencing revealed that the PCR product corresponded to *tubulin alpha 8 like* (*tuba8l*). The *tuba8l* coding sequence was cloned into the EcoRI site of pCS2+. GFP was PCR amplified from the UAS:eGFP-tuba2 plasmid (Asakawa & Kawakami 2010) using primers containing BamHI and ClaI restriction sites (forward primer: 5’- ACGGGATCCGCCACCATGGTGAGCAAGGGCGAGGAGCTG-3’ and reverse primer: 5’- CCGCCGATCGATCTTGTACAGCTCGTCCATGC-3’). Following restriction enzyme digest with BamHI and ClaI, the EGFP coding sequence was cloned in-frame upstream of *tuba8l*.

For transgenesis, *GFP-tuba8l* was excised from pCS2+ using BamHI and XhoI restriction sites with the XhoI end blunted and inserted downstream of the elongation factor 1 promoter in the Tol2 vector pT2KXIGΔin (Urasaki et al. 2006) using the BamHI and ClaI sites with the ClaI end blunted. Tg:(XlEefla1:GFP-tuba8l) transgenic zebrafish were generated using Tol2 transposon-mediated germline transmission (Kotani et al. 2006). Embryos at the 1-cell stage were injected with transposase RNA and pT2KXIGΔin-GFP-tuba8l plasmid and fluorescent embryos were selected at 24 hpf and grown to adulthood. GFP positive embryos from the founder generation were raised to adulthood. The first generation of Tg:(XlEefla1:GFP-tuba8l) were genotyped by crossing to wild type fish and collecting embryos at 24 hpf. Genomic DNA was prepared from approximately 100 embryos per pair and PCR amplification was performed using Taq polymerase (NEB) (forward primer: 5’-ACGGGATCCGCCAC CATGGTGAGCAAGGGCGAGGAG-3’ and reverse primer: 5’- ATGAACTTCAGGGTCAGCTTGC-3’).

### Whole-mount immunohistochemistry

The following primary antibodies (1:500 dilution) were used: rabbit anti-tubulin-detyrosinated (AB3201, EMD Millipore), mouse anti-γ-tubulin clone GTU-88 (T6557, Sigma-Aldrich), and mouse anti-α-tubulin DM1A (T6199, Sigma-Aldrich). The following secondary antibodies were used at 1:1000: goat-anti-rabbit Alexa 488 (A-11008, Invitrogen) and goat-anti-mouse Alexa 488 (A-11001, Invitrogen). Microtubule antibody staining was performed as described (Topczewski and Solnica-Krezel, 1999) with the following modifications: embryos were fixed in 3.7% formaldehyde, 0.2% triton X-100 in microtubule stabilization buffer and fixation time was 1.5 hours at room temperature or overnight at 4°C.

### Imaging

Live and fixed embryos were imaged using a Quorum WAVEFX spinning disk, a Zeiss LSM 510, or a Leica TCS SP8 confocal microscope. Manually dechorionated live embryos were mounted in 0.4-0.8% low melt agarose (Invitrogen) and immunostained embryos were mounted in small drops of 80% glycerol on glass bottom dishes (MatTek).

### Fluorescence intensity measurements of the Dim Zone and PIV analysis

Images were acquired with a Leica TCS SP8 laser scanning confocal microscope using a HC PL APO CS2 20x/0.75 IMM (N.A. 0.75) from Tg:(dclk2DeltaK-GFP) and Tg:(XlEefla1:GFP-tuba81) zebrafish. An oval was fit to the embryo and the region outside of the embryo was masked to exclude irrelevant signal and improve the clarity of the fluorescence profiles. Images were divided along the lateral axis in to 8 equally sized bins that ran the length of the animal - vegetal pole. The mean animal – vegetal intensity profile of each bin was plotted. A central subdivision was used to create the kymographs. Fluorescence intensity profiles from the kymographs were modeled with a polynomial fit and the local minima was used to define the position of the dim zone.

Prior to PIV analysis, stationary background signal was removed by subtracting the mean of all the time points from each time point in the series to better detect changes in intensity. For time lapses of Tg:(dclk2DeltaK-GFP) embryos, analysis was restricted to a region of interest just vegetal to the dim zone and this region of interest was updated for each time point as the dim zone moved vegetally. Images were subdivided into 8x8 pixel interrogation windows with 50% overlaps. For time-lapses of embryos expressing EB3-GFP, analysis was restricted to a rectangular region of interest in the center of the embryo, and images were subdivided into 16x16 pixel interrogation windows with 50% overlap. Each interrogation window was then crosscorrelated with its corresponding window from the next time point to determine the direction and magnitude of fluorescence intensity movement. For the analysis of microtubule flows presented in Figure 2, vectors for each time point were summed over 5 consecutive time points to capture persistent movement and minimize noise. For Figure 2 time steps were: embryo 1: 5.2 min/frame; embryo 2: 6.1 min/frame, embryo 3: 5.1 min/frame; embryo 4: 4.9 min/frame. For the analysis of EB3-GFP comet flows presented in Figure 4, vectors representing movement between individual frames (7.87 sec/frame) were used. The results presented are the mean displacement overtime in the lateral and A-V directions at each time point over the course of similar stages across embryos.

### YSN tracking and identification

e-YSN were identified based on their shape and local absence of fluorescent signal in Tg:(dclk2DeltaK-GFP) and Tg:(XlEefla1:GFP-tuba81) time-lapse movies and tracked using the ImageJ plugin “Manual Tracking”. e-YSN were identified in H2A-GFP expressed embryos based on shape and position within the Z-stack.

## Acknowledgements

AB thanks R. Winklbauer for many helpful discussions and insightful comments on the manuscript. We thank A. Akhmanova, M. Mione and K. Sampath for reagents and Henry Hong for confocal assistance.

---

**Supplementary Table 1:**
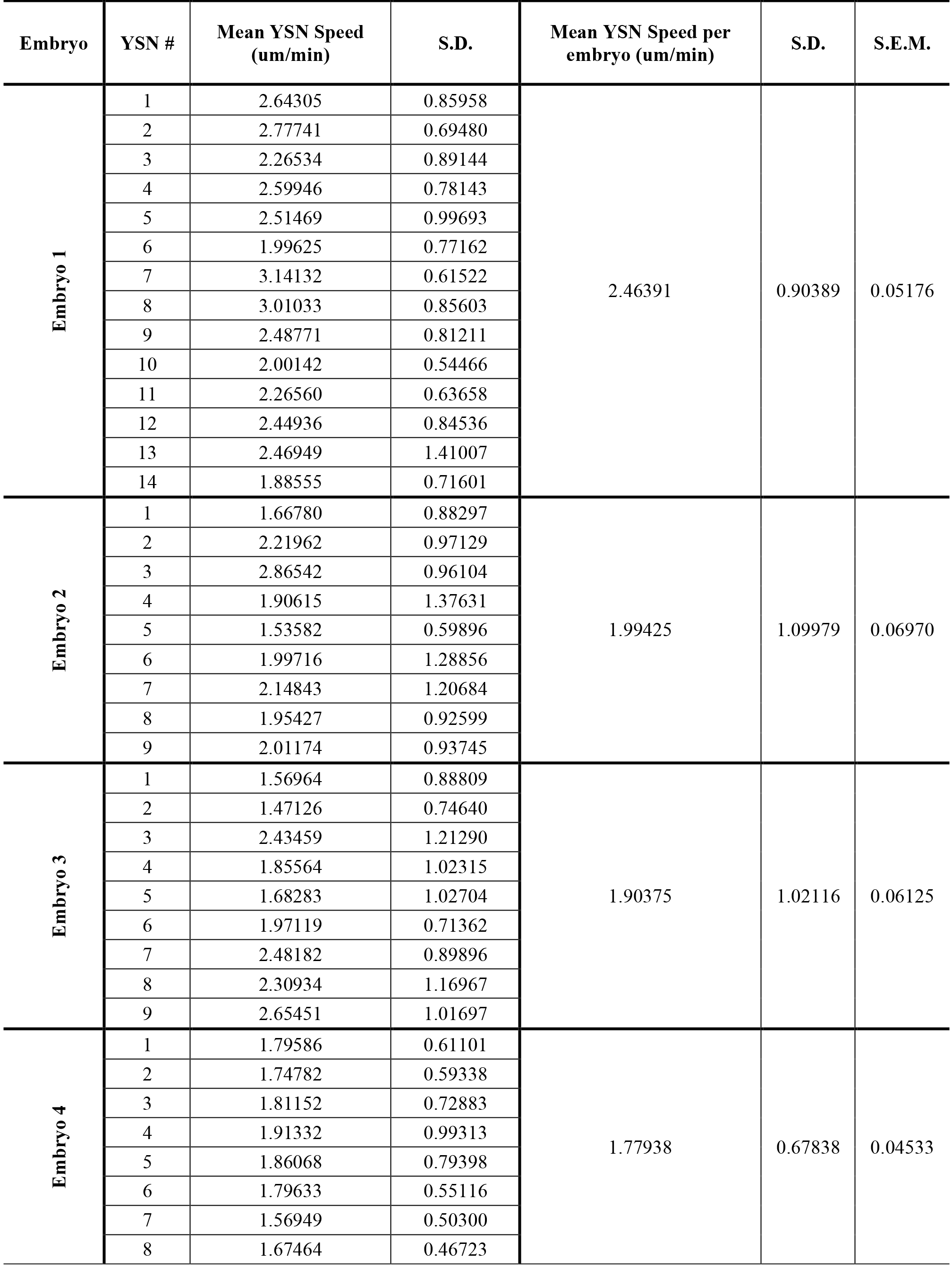
Individual mean e-YSN speeds.

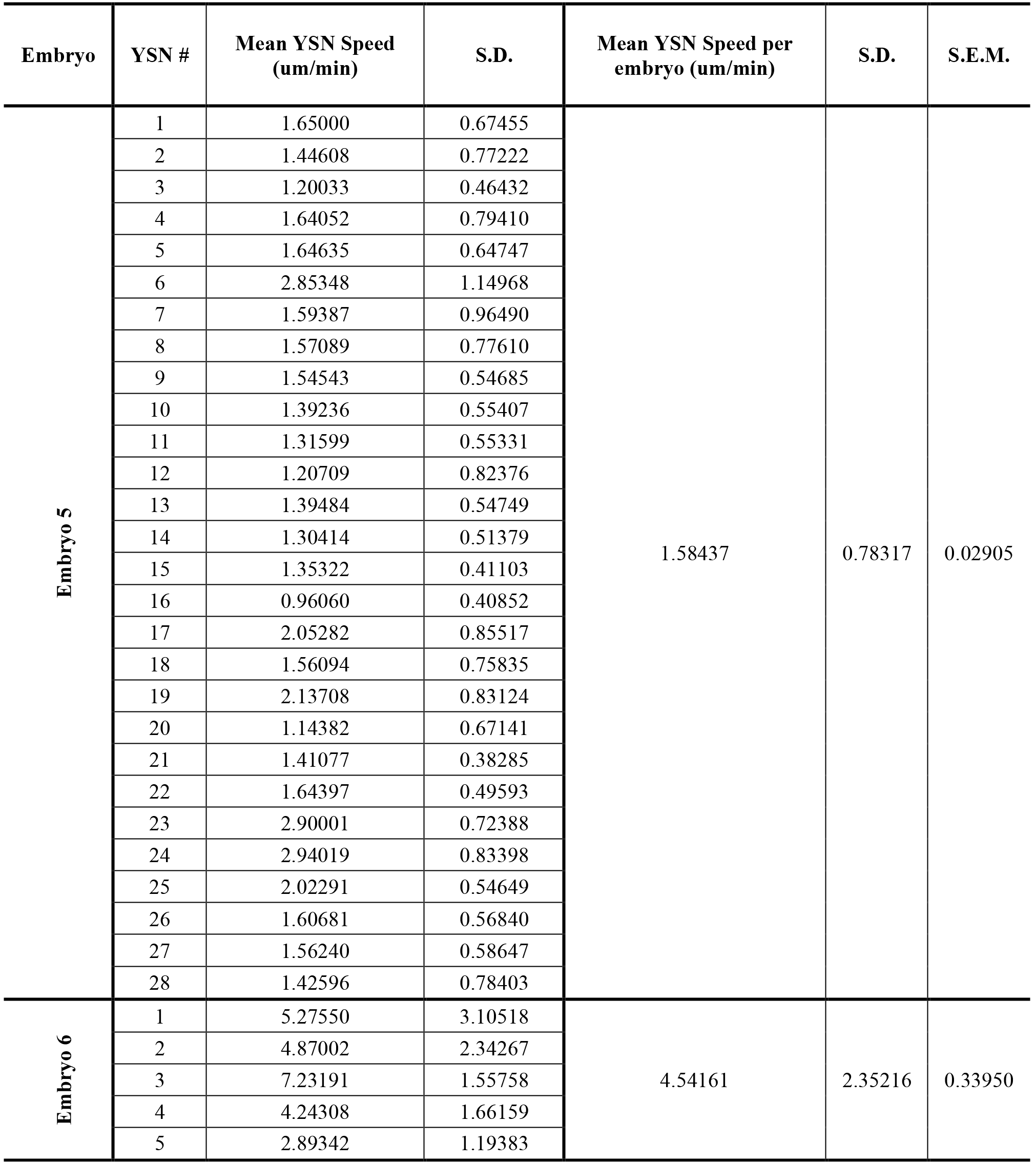

## Supplementary Figure Legends

### Movie 1

Confocal time-lapse movie of Tg(XlEef1a1:dclk2DeltaK-GFP) embryo from sphere stage to 80% epiboly. Lateral view with the animal pole to the top.

### Movie 2

Confocal time-lapse movie of embryo expressing EB3-GFP. Lateral view with animal pole to the top.

### Movie 3

Confocal time-lapse movie of embryo expressing EB3-GFP. e-YSN can be seen emerging from regions where EB3-GFP comets are emanating (arrows). Lateral view, animal pole towards left. Bright region at the top is the blastoderm.

### Movie 4

Spinning disk confocal time-lapse movie of Tg (XlEef1a:eGFP-tubα8l) embryo with H2A-GFP labeled e-YSN. Lateral view with animal pole towards the top.

### Movie 5

Confocal time-lapse movie of H2A-GFP expressing control embryo. Lateral view with animal pole towards the upper right.

### Movie 6

Confocal time-lapse movie of H2A-GFP and C-Syne2a expressing embryo. Lateral view with animal pole towards the upper left.

